# A decision-theoretic model of multistability: perceptual switches as internal actions

**DOI:** 10.1101/2024.12.06.627286

**Authors:** Shervin Safavi, Peter Dayan

## Abstract

Perceptual multistability has been studied for centuries using a diverse collection of approaches. Insights derived from this phenomenon range from core principles of information processing, such as perceptual inference, to high-level concerns, such as visual awareness. The dominant computational explanations of perceptual multistability are based on the Helmholtzian view of perception as inverse inference. However, these approaches struggle to account for the crucial role played by value, e.g., with percepts paired with reward dominating for longer periods than unpaired ones. In this study, we formulate perceptual multistability in terms of dynamic, value-based, choice, employing the formalism of a partially observable Markov decision process (POMDP). We use binocular rivalry as an example, considering different explicit and implicit sources of reward (and punishment) for each percept. The resulting values are time-dependent and influenced by novelty as a form of exploration. The solution of the POMDP is the optimal perceptual policy, and we show that this can replicate and explain several characteristics of binocular rivalry, ranging from classic hallmarks such as apparently spontaneous random switches with approximately gamma-distributed dominance periods to more subtle aspects such as the rich temporal dynamics of perceptual switching rates. Overall, our decision-theoretic perspective on perceptual multistability not only accounts for a wealth of unexplained data, but also opens up modern conceptions of internal reinforcement learning in service of understanding perceptual phenomena, and sensory processing more generally.

## Introduction

Multistable perception is the perceptual phenomenon of dynamic alternation that arises when a single sensory input has more than one interpretation or explanation. This phenomenon has been studied for decades (Leopold et al., 1999; Blake et al., 2002; Brascamp, Klink, et al., 2015; Brascamp, Sterzer, et al., 2018; Safavi and Dayan, 2022), across a variety of species, including Drosophila (Toepfer et al., 2018), fish and reptiles (Carter et al., 2020), mice (Palagina et al., 2017) primates (Blake et al., 2002), and human (Zaretskaya, 2021), and almost all sensory modalities (Schwartz et al., 2012; Alais, Keys, et al., 2021). Studying the phenomenon has provided insights into a diverse set of cognitive functions, including, attention (Watanabe et al., 2011; Maier et al., 2021), visual awareness (Seth et al., 2022; Panagiotaropoulos, 2024), perceptual inference (Brascamp, Sterzer, et al., 2018; Hardstone et al., 2021), perceptual decision-making (Krug, 2020), emotion processing (Bannerman et al., 2011), and more recently, even interoception (Praag et al., 2024; Veillette et al., 2024) and language (Styrnal et al., 2023). Moreover, various aspects of perceptual multistability have been found to differ in a range of psychiatric conditions (see, e. g., Ye et al., 2019; Jia et al., 2020).

The traditional account of multistability is based on a passive account of perception in which it constitutes under-controlled inference about the contents of the world (Dayan, 1998; Schrater et al., 2007; Sundareswara et al., 2008; Hohwy et al., 2008a; Reichert et al., 2011; Moreno-Bote et al., 2011; Gershman et al., 2014; Weilnhammer et al., 2017; Leptourgos et al., 2020; Cao et al., 2021; Weilnhammer, Chikermane, et al., 2021) — also see Brascamp, Sterzer, et al. (2018), for a review. However, it is difficult to reconcile this formulation with a large body of recent data concerning both behaviour and the neural substrate of perceptual multistablity (Martin et al., 2021) which collectively imply that fundamental concepts such as reward (and broadly value) and choice are missing (for a review see, Safavi and Dayan, 2022). For instance, when a percept is paired with reward, it dominates for longer periods than if it is not; and, equally, multiple brain systems, such as the prefrontal cortex (Michel, 2022; Brascamp, Sterzer, et al., 2018; Safavi, Kapoor, et al., 2014) and the anterior cingulate cortex (Lyu et al., 2022; Drew et al., 2021; Gelbard-Sagiv et al., 2018), that are generally engaged in value-based operations such as cognitive control and decision-making are also involved in multistable perception. Certainly, areas of the brain are far from singular in their computational roles; however, reconciling the neural and behavioural data in light of a decision-theoretic approach to perceptual multistability could be more plausible compared to traditional value-free accounts.

We formulate perceptual multistability as a dynamic, value-based decision-making process, ex-tending the traditional account via the formalism of a partially observable Markov decision process (POMDP). As in conventional Bayesian accounts of perception, we consider there to be a single actual state of the world at any one time that must be inferred from sensory observations; but (in the simplest case), for which there are two plausible, but incompatible candidates (Dayan, 1998; Brascamp, Sterzer, et al., 2018). Each percept is potentially associated with different sources of rewards or punishments, and switching between percepts, more precisely perceptual selection, is a form of (costly) *internal action* (Dayan, 2012) – the attentional equivalent of the external action of moving eye gaze from one object to another. This makes perception arise as a choice rather than a passive inference. The approximate solution of the POMDP is the optimal perceptual policy, which we show to replicate and explain several aspects of rivalry.

Based on a diverse set of behavioral data, we demonstrate that the new formulation not only explains key characteristics of multistability (Levelt, 1965; Blake et al., 2002; Brascamp, Klink, et al., 2015) but also accounts for several elusive aspects that previous frameworks struggle to explain (Balcetis et al., 2012; Wilbertz, van Slooten, et al., 2014; Marx et al., 2015; Wilbertz, van Kemenade, et al., 2017; Pothast et al., 2024; Lunghi et al., 2023; Lunghi et al., 2019; Haas, 2021; Haas, 2021; Suzuki et al., 2007; Dieter et al., 2016). The latter includes reward (Balcetis et al., 2012; Wilbertz, van Slooten, et al., 2014; Marx et al., 2015; Wilbertz, van Kemenade, et al., 2017; Pothast et al., 2024; Lunghi et al., 2023; Lunghi et al., 2019; Haas, 2021; Haas, 2021) and task (Dieter et al., 2016) modulation of perception during multistablity, as well as the rich temporal dynamics of perception across three distinct timescales (Suzuki et al., 2007). Factors that allow us to account for such a diverse set of observations are not only due to the dynamic and decision-theoretic nature of our proposed model but also from the richness of reward function – most notably incorporating the recent computational models of aesthetic reward in our framework.

Our treatment of perceptual multistability is grounded in the value awarded to a pure sensory process that its dynamics shape the dynamics of perception during multistablity. Aesthetic aspects of an image can, in fact, bias perceptual preference in perceptual multistability (Huang et al., 2013; Mo et al., 2016; Shang et al., 2020; Tsikandilakis et al., 2019). Even in the frequently used case of rivalry between face and house images, faces, which occupy a special position in our aesthetic lives (Rhodes, 2006), have longer dominance times (Persike et al., 2014). Additionally, the relationship between complexity and aesthetics (Nath et al., 2024) implies complex stimuli also dominate for longer, which is experimentally confirmed (Rogers et al., 1977; Alais and Melcher, 2007; Daini et al., 2010). Most importantly, a particular characteristic of aesthetic value is that it changes over the course of observation, captured, for instance, in modern theories (e.g., Brielmann and Dayan, 2022). We therefore also *quantitatively* examine the extent to which the temporal dynamics of multistability could be inherited from the temporal dynamics of aesthetics. Furthermore, along with other cognitive factors, aesthetic value, with its evident ethological roots (Rhodes, 2006; Chittka et al., 2006; Iigaya, Yi, Wahle, Tanwisuth, et al., 2021), is a key component of the reward associated with a percept and thus the decision-theoretic reason to perceive it (Scocchia et al., 2014; Safavi and Dayan, 2022).

Lastly, along with these theoretical and quantitative analyses of the multistablity behaviours, we also suggest a range of empirical consequences of our account for the neural side of multistablity. For instance, we consider the implications for the involvement of value- and policy-related brain circuits such as the anterior cingulate cortex (ACC, Lumer et al., 1998; Gelbard-Sagiv et al., 2018; Drew et al., 2021; Lyu et al., 2022), pulvinar (Wilke et al., 2009), locus coeruleus (LC, see, e. g., Einhauser et al., 2008), as well as (the controversial) involvement of prefrontal cortex (PFC, Zaretskaya and Narinyan, 2014; Safavi, Kapoor, et al., 2014; Tsuchiya, Wilke, et al., 2015; Odegaard et al., 2017; Block, 2019; Michel, 2022; Canales-Johnson et al., 2023), which are more conventionally tied to value-dependent operations such as cognitive control and foraging (Montague et al., 2002; Niv, 2009; Lee et al., 2012; Monosov et al., 2020; Holroyd et al., 2021a; Rudebeck et al., 2022). Notably, this also includes the neural systems that have been suggested to compute aesthetic value, namely vlPFC (Iigaya, Yi, Wahle, Tanwisuth, et al., 2023) and ACC (Cloutier et al., 2008) which are also involved in perceptual multistability.

In sum, our decision-theoretic approach connects classical approaches to perception with modern ideas from reinforcement learning. Our approach, not only synergizes with Helmholtzian account of perceptual multistability (Dayan, 1998; Schrater et al., 2007; Sundareswara et al., 2008; Hohwy et al., 2008a; Reichert et al., 2011; Moreno-Bote et al., 2011; Gershman et al., 2014; Weilnhammer et al., 2017; Leptourgos et al., 2020; Cao et al., 2021) and offers a more comprehensive treatment of computational and algorithmic facets of perceptual multistability, but also it links new ideas about the genesis and effect of value with modern conceptions about internal reinforcement learning (Hazy et al., 2007; Dayan, 2012; Lieder et al., 2018) and thereby helps explain one of the most venerable and well-studied phenomena in sensory processing.

## Results

We model the phenomenon of binocular rivalry, which is one of the most extensively studied forms of perceptual multistability (Blake et al., 2002). In the canonical form of binocular rivalry, a participant’s two eyes are shown two distinct images (***Figure 1A***) over an extended period. This leads to spontaneous switches between two percepts, typically, each associated with one image. The image currently perceived is referred to as being dominant (***Figure 1B***); the periods of dominance more (Pastukhov et al., 2013), or less (Brascamp, Ee, et al., 2005), follow a gamma-like distribution (***Figure 1C***). There are also piecemeal percepts, which involve elements of both images, but, for simplicity, we do not consider them in our model.

**Figure 1.**
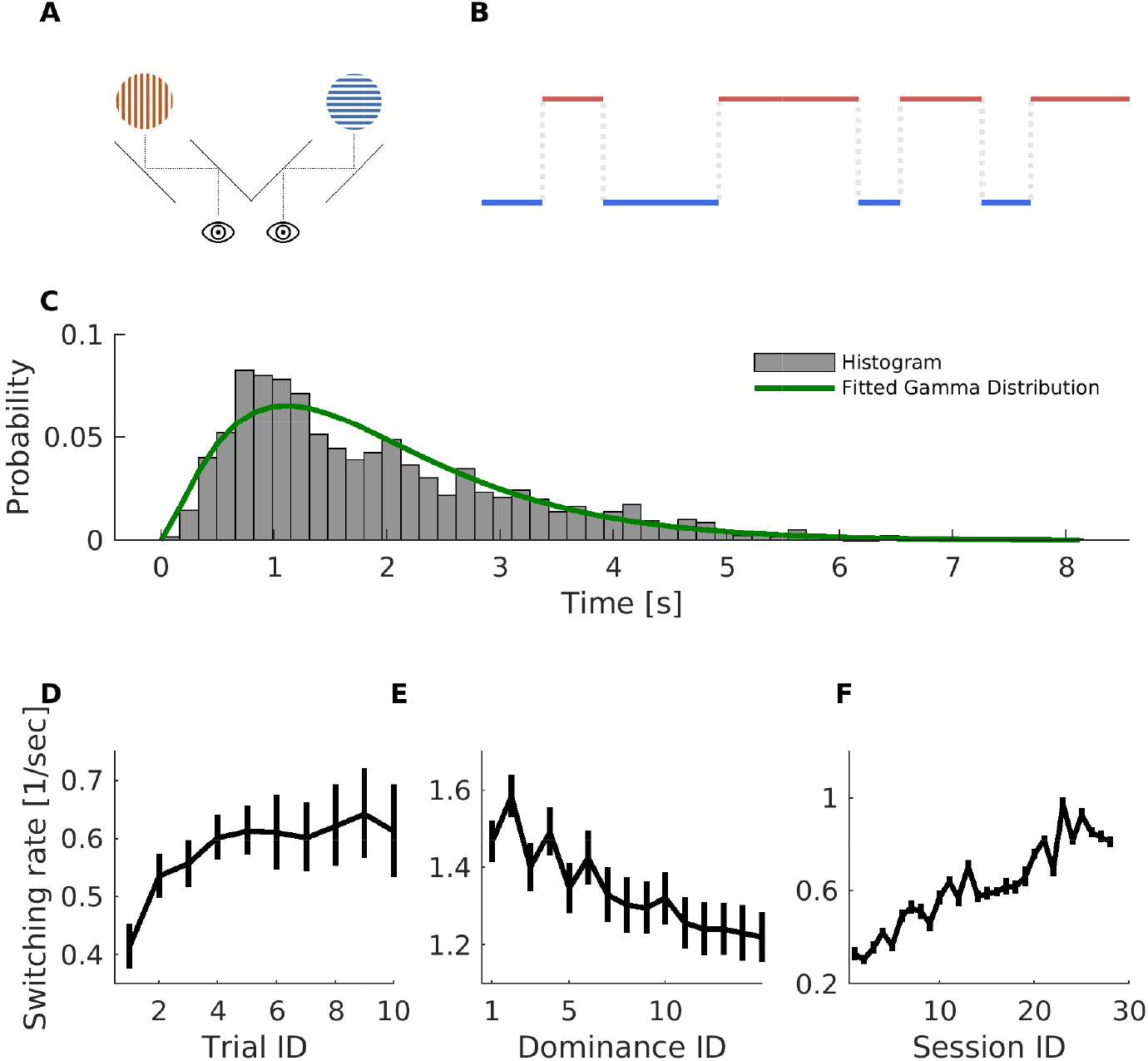
Basics of binocular rivalry. **(A)** Binocular rivalry involves a participant’s two eyes seeing images that differ by more than just depth disparity. **(B)** Participants typically perceive one of the presented images, vertical (left, red) or horizontal (right, blue) gratings with *dominance periods* (when either the left or right image is exclusively perceived), and *perceptual switches* (transition from one percept to the other). **(C)** Dominance durations in binocular rivalry share the common statistical property of being approximately gamma-like distributed. The example distribution is based on data presented in Kapoor et al. (2020). **(D)** The initial increase of switching rate over 10 trials, (including all dominance periods within a trial) and averaged across 40 participants for the first 5 trials, and 16 participants for the last 5 trials (resulting in larger error bars for trials 6 through 10; see, Suzuki et al., 2007, for more details). **(E)** Decreases in switching rate across dominance instances within individual trials (each data point is an average across 21 sessions measured for one participant). **(F)** Long-term speed-up of switching rate across sessions for one participant (individual data point for each session is average across all dominance periods in all 20 trials of each session, resulting in small error bars). In D-E, error bars are mean ± SEM, and data adopted from Suzuki et al. (2007).

Perceptual multistability has rich temporal dynamics. Suzuki et al. (2007) reported that the rate of perceptual switches changes on three distinct timescales: naive participants experience an initial increase in the rate of switches over the course of dozens of trials (***Figure 1D***; a similar observation was made by Brown, 1955). Then, over the course of a single trial (tens of seconds), participants experience a gradual decrease in the rate of perceptual switches (***Figure 1E***; van Ee, 2005, also reported a somewhat similar observation), Finally, the switching rate slowly increases across several sessions (***Figure 1F***; Weilnhammer, Fritsch, et al., 2021a, also reported a similar slow dynamics). Using our model, we replicate and explain these rich dynamics along with several other characteristics of perceptual multistablity.

### Modelling binocular rivalry with a POMDP

We approach perceptual selection as an internal action, and dominance sequences as decision sequences. Thus, we model binocular rivalry (see ***Figure 2*** for a schematic) based on a well-established model of decision-making — the partially observable Markov decision process (POMDP, Littman, 2009, and see Methods section, for more details). Based on the conventional Bayesian accounts of perception (Yuille et al., 2006; Dayan, 1998; Brascamp, Sterzer, et al., 2018), we consider there to be a single actual state of the world at any one time that must be inferred from sensory observations. Given that, there are two input images, we assume that there are two plausible but incompatible candidates for this inference. However, Bayesian perception ends at the estimation of the posterior distribution over these two candidates; we take the further Bayesian decision-theoretic step *of considering an explicit choice between the candidates, treating this as an internal decision or action of perceptual selection* (Dayan, 2012). This action is accompanied by reduced observation noise (see section Sensory, perceptual and decision processes), which, amongst other consequences, entails a stronger belief about the perceived state (dominant percept). The action is also endowed with sources of rewards or punishments, which are themselves dynamic. For instance, each image has an aesthetic value (how much we like it), which changes over time (e.g., as we get bored after multiple exposures; Brielmann, Berentelg, et al., 2023).

**Figure 2.**
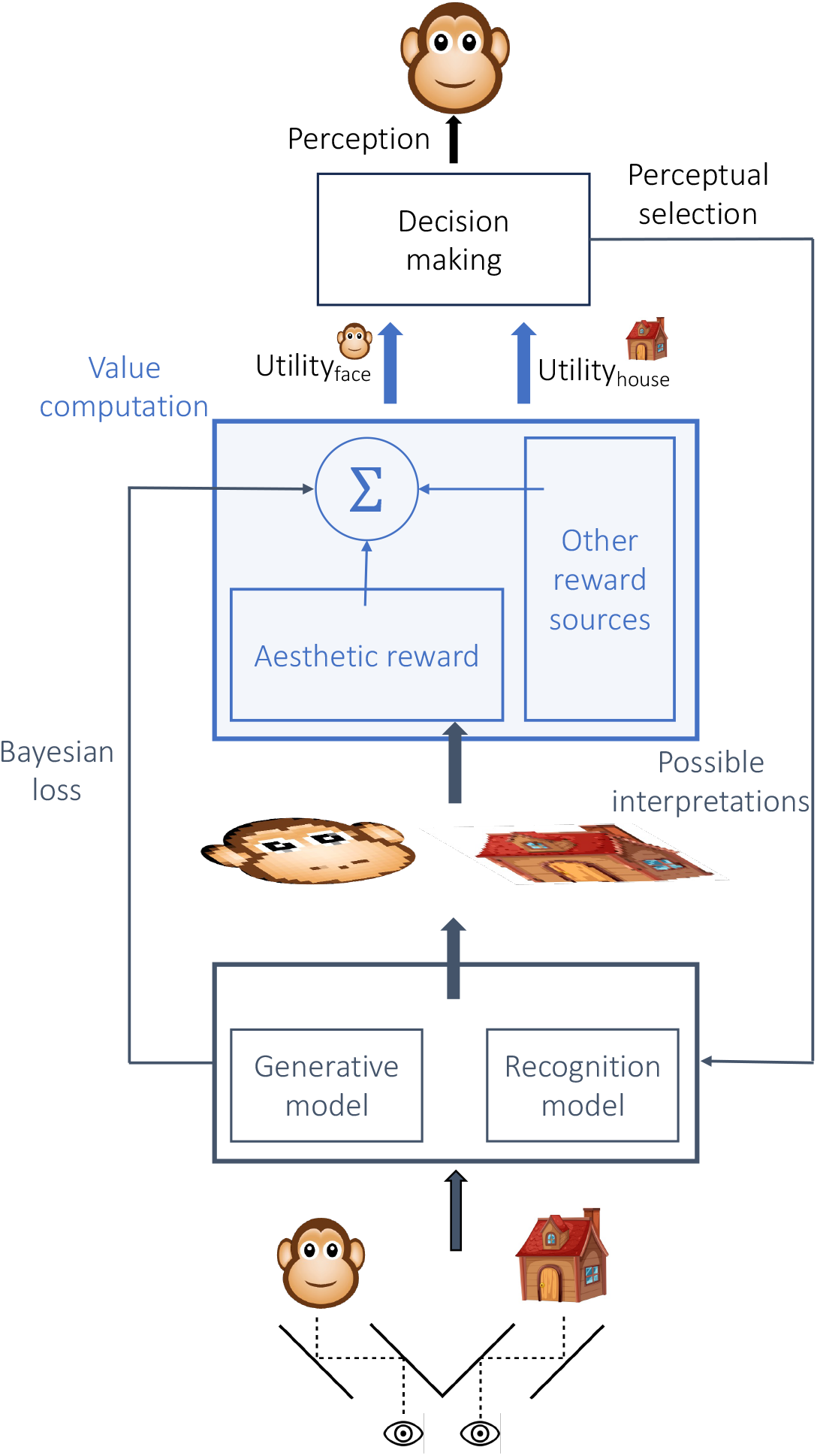
Schematic demonstration of the decision-theoretic model. In our model, the agent receives two sensory inputs (one per eye). As in conventional Bayesian accounts of perception, we assume our agent has a generative and recognition model that leads to two possible interpretations of sensory inputs (also affected by noise). In the next stage, each possible interpretation of sensory input goes through a *valuation* process that determines the utility of each perception and how it evolves over time. The utility of each perception depends on various factors, such as associated reward (e.g., external and aesthetic reward), error of the generative model in explaining the input, and other factors. Perceptual action is the selection of a perception with a larger utility.

The agent needs to take into account all contributing factors (the momentary total utility of each possible percept, belief strength, etc) to decide which percept should dominate. These factors include conventional components from classical Helmholtzian treatments to perceptual multistability, as well as components originating in our value-centred conceptualisation. An example conventional component is the conventional loss that purely inference-based models of perceptual multistability emphasize (Dayan, 1998; Schrater et al., 2007; Sundareswara et al., 2008; Hohwy et al., 2008a; Reichert et al., 2011; Moreno-Bote et al., 2011; Gershman et al., 2014; Weilnhammer et al., 2017; Leptourgos et al., 2020; Cao et al., 2021), which counts the unlikelihood of the agent’s observations given the current decision of the agent about the state of the world. An example of value-centered component is an approximate form of the value of information that is associated with a stimulus which is incompletely known (Dayan and Sejnowski, 1996, also see section Sensory, perceptual and decision processes, for more details).

The policy of the agent – i.e., the systematic (and potentially probabilistic) mapping of the images and the history of its past perceptual choices to its next sensory decision – is the solution of the POMDP. This means, finding the optimal actions, i.e., selecting actions that lead to the maximum utility for the agent (see section Partially observable Markov decision process, for more details). We show that this policy replicates, and thereby explains, several aspects of rivalry (as discussed in the next sections).

### Simplified POMDP and multistability hallmarks

We start from a version of our POMDP (described in Methods) in which the reward function is simplified. This can already capture some key hallmarks of multistability. Perhaps the simplest contributor to reward is the aesthetic value of a percept (Brielmann and Dayan, 2022). This should be present across a wide range of experimental settings (since participants are always exposed to some form of sensory input). For our simplified model, we consider this value to evolve according to a simple, phenomenological, case of exponential decay. The rationale behind this choice is that aesthetic value wanes with boredom and/or fatigue – at least for the simple stimuli that are often used in binocular rivalry experiments (such as grating patterns).

Given this exponentially decaying reward and belief fluctuations, the agent spontaneously switches between percepts, leading to a variety of dominance periods (***Figure 3A***). The randomness in the switches comes from belief fluctuations rooted in observation noise. More precisely, when the aesthetic value decays sufficiently, fluctuations induced by the observation noise can lead to a switch (***Figure 3B***). Thus, there is a mean dominance period that depends on the characteristic decay of phenomenological aesthetic value, with variability stemming from observation noise (***Figure 3A-B***). Interestingly, spontaneous dominance periods are also (approximately) gamma-distributed (***Figure 3C***, green curve) as also observed in humans and monkeys (Blake et al., 2002). This is also preserved under further manipulations (see e.g., Supplementary Figure 1, for a setting with asymmetric reward).

**Figure 3.**
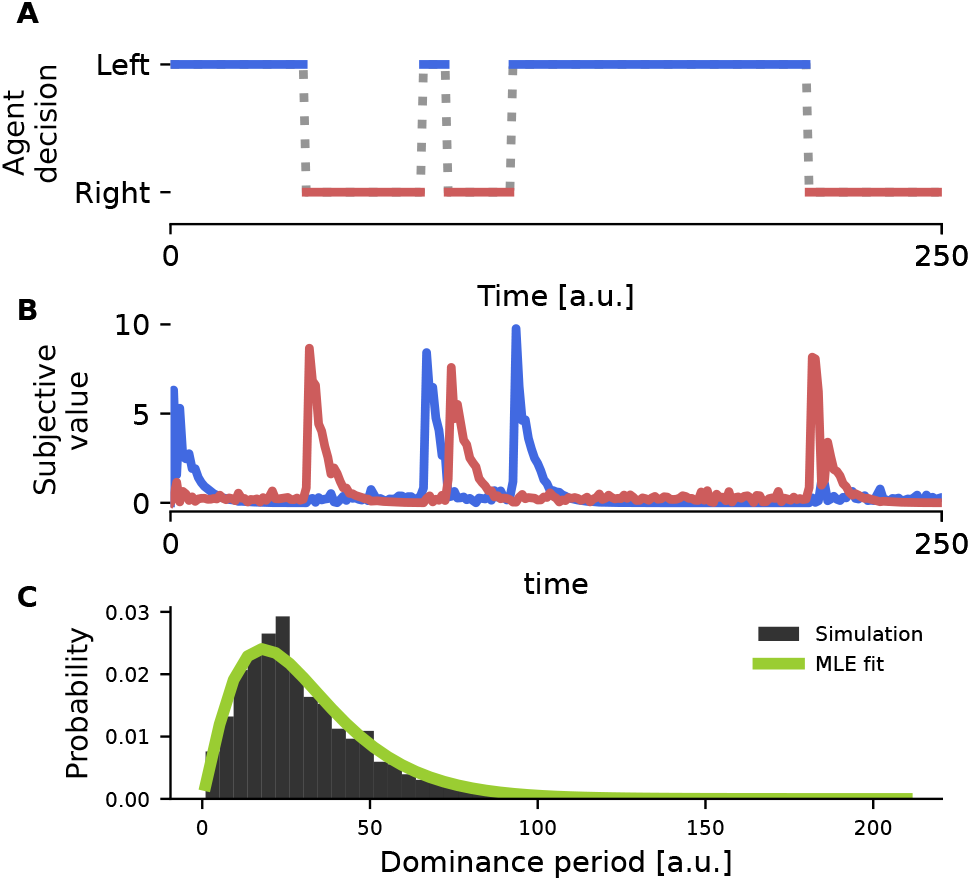
A simple POMDP captures the statistical hallmarks of perceptual multistability. **(A)** An example trajectory of the agent’s decision sequence between left and right images in a binocular rivalry task. **(B)** Reward function of the simplified POMDP, which is a phenomenological aesthetic value modelled with exponential decay, which fluctuates due to observation noise. **(C)** Distribution of dominance periods for an extended length of simulation. Gamma distribution fitted with maximum likelihood estimation (MLE) and indicated with a green line.

The design of this simple model resonates with the key features of classical biophysical models of binocular rivalry, where noise and adaptation are the key factors shaping the dynamics. Our formulation offers a functional interpretation for traditional biophysical characteristics such as fatigue, e.g., putting such biophysical constructs next to other cognitive factors modulating perception (Scocchia et al., 2014; Safavi and Dayan, 2022). Notably, as discussed later (section Perceptual privileges through reward asymmetries and Levelt’s propositions), this formulation allows us to explain the essential aspects of perceptual multistability such as Levelt’s propositions (Levelt, 1965), and modern conceptualization such as reward modulation of perception (Balcetis et al., 2012; Wilbertz, van Slooten, et al., 2014; Marx et al., 2015; Wilbertz, van Kemenade, et al., 2017), based on a common computational principle in a single framework (that is inspired by reinforcement learning).

### Perceptual events are orchestrated by dynamics of the competing value

To explain a broader range of observations in multistablity we incorporate a more complex reward function in the model. Although the simple decision-making model of ***Figure 3*** can capture hallmarks of binocular rivalry (apparently spontaneous switches and approximately gamma-distributed dominance periods), the phenomenological account of aesthetic value does not have sufficient capacity to capture all the intricate aspects of multistability. Thus, in the following, we employ a more complex model of aesthetic value and demonstrate how it shapes the dynamic of perceptual switches.

A recent theory of the aesthetic value of sensory objects (Brielmann and Dayan, 2022) has suggested that, aesthetic value can be derived from the task of optimising a sensory system to process present and future stimuli well. In this framework, an agent is assumed to have a momentary or contextual probabilistic model of what it expects to sense now (called the ‘system state’) and a more general probabilistic model for the sensory input it expects in the long run (the ‘expected true distribution’). Perceiving a stimulus (or in the case of binocular rivalry, one of the two stimuli) leads to a mandatory adaptation or learning of the system state. This then leads to two underlying components for the aesthetic value: a) the fluency with which it can be processed (i.e., how well it can be encompassed by the system state); and b) the effect of plasticity occasioned by perceiving this stimulus for the fluency with which likely future stimuli will be processed. As the system state moves, the values of the two components change, and so the aesthetic value changes too (described in more detail in Model of aesthetic value).

Within this model of aesthetic value, we make the substantial simplification of representing stimuli (and the agent’s estimate of the stimuli) in a two-dimensional feature space. In binocular rivalry, two stimuli are available to the agent, thus they are characterised by two distinct coordinates in this feature space (***Figure 4A***, blue and red circles). The initial location of the system state’s mean (***Figure 4A***, black circle) is chosen such that it is equidistant from the location of both stimuli (which implies that the stimuli have equal aesthetic value).

**Figure 4.**
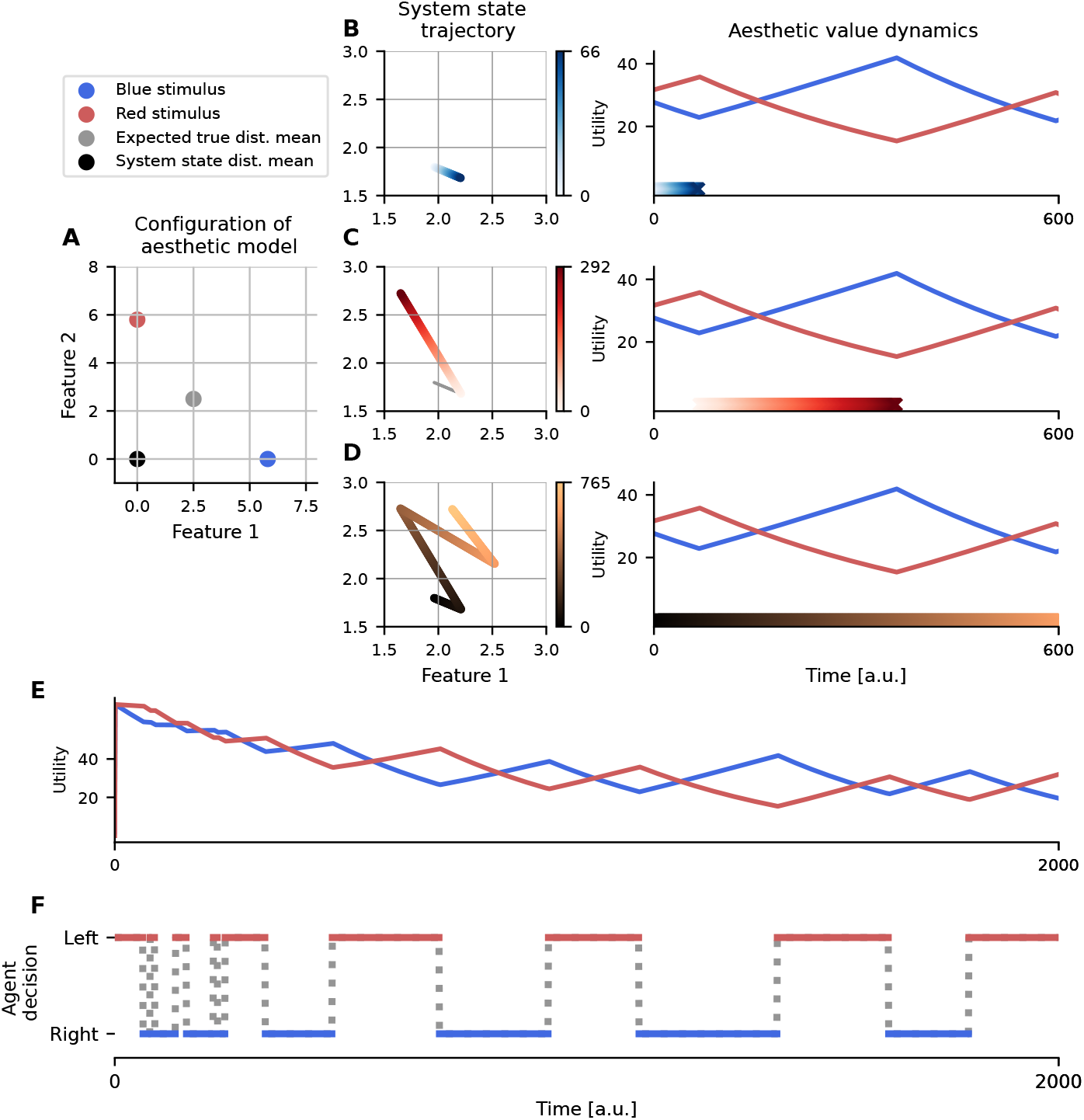
Push-pull mechanism of perceptual alternations governed by dynamics of value. **(A)** Representation of two rivalling stimuli (red and blue circles) and the initial location of the system state mean and expected true distribution mean in the model of aesthetic value (black and grey circle respectively). **(B)** When the blue stimulus is the dominant percept, the system state moves toward its representation in the feature space (left figure) and the aesthetic value of the blue stimuli decays (blue trace in the right figure). Simultaneously, the system state gets further from the red stimulus, which leads to an increase in aesthetic value for the red stimulus if it becomes dominant (red trace in the right figure). Temporal evolution is colour-coded in both left and right figures according to the gradient colourbar in the middle. **(C-D)** Similar to (B), but for the second perceptual switch (C) and both switches together (D). **(E)** Global dynamics of the aesthetic reward for red and blue stimuli during an extended period of binocular stimulation. **(F)** Similar to ***Figure 3A***, an exemplary trajectory of the agent’s decision sequence temporally aligned with (E) during the binocular rivalry task.

The agent enjoys the dynamically changing aesthetic reward of the stimulus that is perceived at each moment. For instance, when the agent perceives the blue stimulus, it starts consuming its aesthetic reward. In the course of the perception of the blue stimulus, the system state gradually evolves toward the location of the blue stimulus in the feature space. The change in the location of the system state in relation to the blue stimulus and the expected true distribution determines how the aesthetic value changes over time. As the system state moves toward the blue stimulus (gets closer), the agent better accounts for and processes the blue stimulus (fluency component of the aesthetic value), however, the dynamism of the system state dwindles (learning component of the aesthetic value). This implies that the learning component of the aesthetic reward for the blue stimulus decreases over the period that the agent perceives this stimulus, and it implies that the aesthetic value (which is the sum of both the fluency and the learning components) of a dominant stimulus decreases over time (***Figure 4B***). This creates a *pushing* for a perceptual switch. At the same time, the system state gets further from the red stimulus, which also changes the aesthetic reward that the agent *could accrue* from the red stimulus *if* it were switched into. Thus, the agent continuously estimates the (counterfactual) aesthetic reward of the alternative perception (red stimulus in this case). Given that the system state is far and getting even further from the red stimulus, the (counterfactual) evolution of the system state toward the red stimulus would be large (see Model of aesthetic value, ***Equation 11***). In terms of the formulation of aesthetic value we use, it means that perception of the red stimulus would generate substantial learning, or in other words, this change has a high long-term utility for the agent (see Model of aesthetic value, and in particular ***Equation 14***). Thus, in the course of the perception of the currently dominant stimulus, i.e., closure of the system state to the dominant perception and distance from the suppressed one, the (counterfactual) aesthetic reward of the suppressed (red stimulus) gradually increases (due to learning component of aesthetic value). This is therefore *pulling* for a perceptual switch. At some point, more rewards would be available by switching to the alternative percept. These together create a periodic (yet random, due to belief fluctuations) push-pull mechanism, and thus perceptual alternations (***Figure 4D***).

Our proposed push-pull mechanism for perceptual switching also resonates with dynamic systems models of perceptual multistability (e. g., Brascamp, van Ee, et al., 2006). In these models, the adaptation of the dominant percept may correspond to a push factor, and recovery from that adaptation may act as a pull factor, particularly noting the gradual nature of perceptual switches described in these models. However, the pull factor in our model can be cognitively richer. For instance, in continuous flash suppression (CFS; see Tsuchiya and Koch, 2004, a similar phenomenon to binocular rivalry, where a contentful image presented to one eye is suppressed by dynamic patterns presented to the other eye), faces with fearful expressions have shorter suppression periods compared to neutral or happy expressions. In such cases, it has been suggested that fearful faces are privileged because of their emotional content (Yang et al., 2007; Sterzer et al., 2011), which, in the context of our framework, can be considered a stronger pulling force by the suppressed percept for switching (or, more precisely for CFS, breaking through awareness). The same applies to other forms of cognitive associations (e. g., for the association of external reward, see Lunghi et al., 2023).

Overall, our results suggest that the dynamics of perception are shaped (at least in part) by the competing and dynamic value of sensory stimuli in binocular rivalry. Notably, this relies not only on the value of the perceived stimuli, but also, on the value of the alternative perception and its dynamics (notably that can be cognitively rich, e.g., through affective charges). This provides an *active* perspective compared to classical biophysical and normative models where switches are driven primarily by (manually included) adaptation/fatigue and noise, i.e., a push due to the cessation of the currently dominant percept, and recovery from adaptation as the pull factor.

### Perceptual privileges through reward asymmetries and Levelt’s propositions

As mentioned earlier, multiple sensory and cognitive factors can favour particular percepts in the context of perceptual multistablity. We consider asymmetries in reward function by manipulating different components of this function (e.g., differential aesthetic preferences) and demonstrate it can explain several established aspects of perceptual multistability.

We start with the instrumental imperative to execute actions associated with an aspect of the percept (e. g., to avoid a negative outcome or gain a positive one). In the simplest setting where no instrumental reward is involved (***Figure 5Ai***, resembling the most conventional binocular rivalry experiments), we observe equi-dominance (***Figure 5Bi***, as seen in previous studies, Moreno-Bote et al., 2010). However, consider the case where there is an explicitly learned association between percept and reward. This can institute an asymmetry in the rewards associated with perceiving a stimulus, so that either the left (***Figure 5Aii***) right (***Figure 5Aiii***) images have a higher value (as empirically tested in, Wilbertz, van Slooten, et al., 2014; Marx et al., 2015), leading to increased dominance of the corresponding perception (***Figure 5 Bii*** and Biii, Wilcoxon rank-sum test, *p* < 10^−20^).

**Figure 5.**
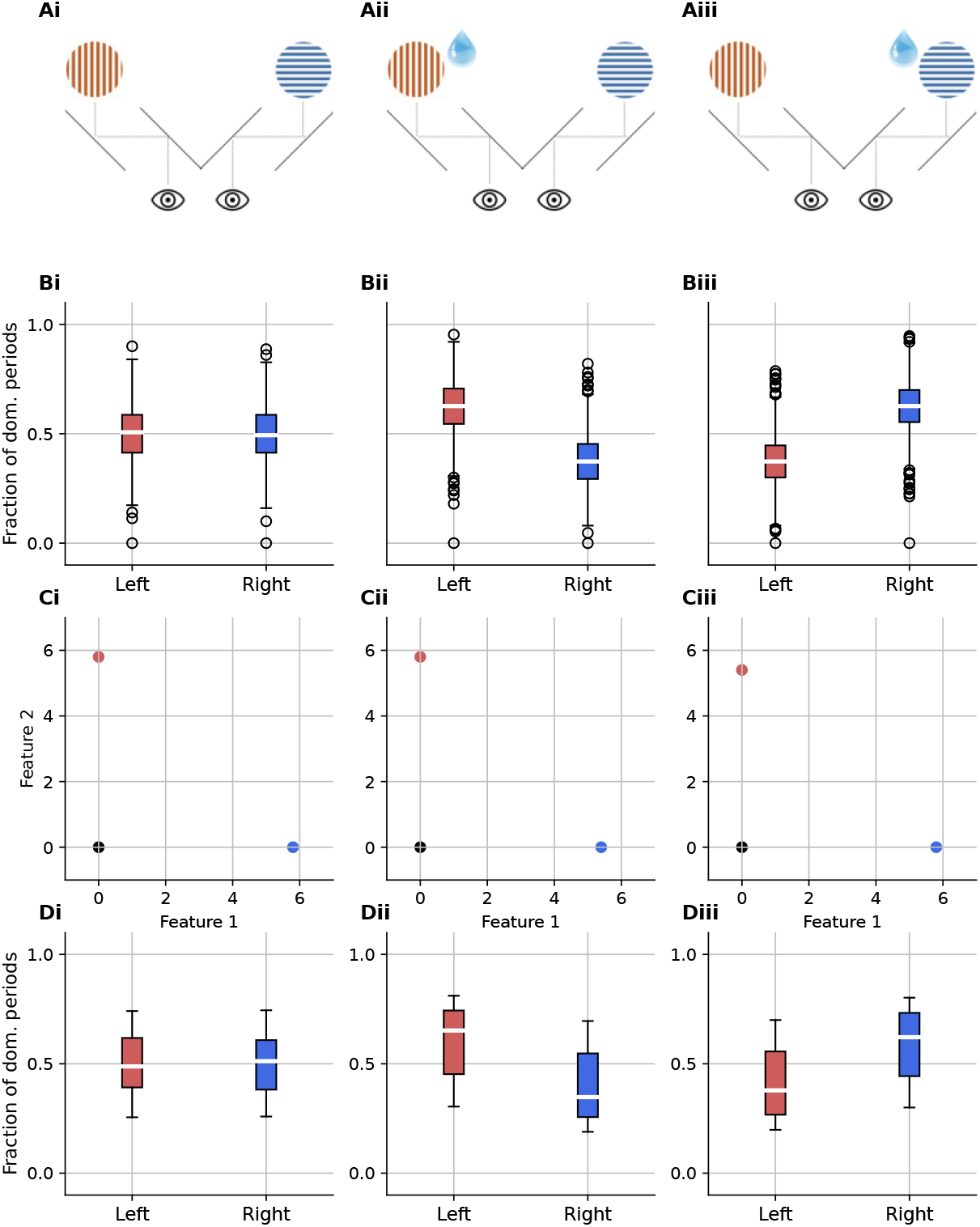
Modulation of dominance periods by different sources of reward. **(Ai-Aiii)** Reward structure of the binocular simulations: (Ai) no reward asymmetry, (Aii) reward associated with the left stimulus, and (Aiii) right stimulus. **(Bi-Biii)** Distribution of dominance periods (as a fraction of the entire trial) for left and right stimuli in different reward structures corresponding to Ai-Aiii. **(Ci-Ciii)** Different geometry of parameters in the aesthetic model to create similar (symmetric or asymmetric) reward structures as in Ai-Aiii. Each panel represents the location of two stimuli of binocular rivalry (red and blue circles) and the mean of the system state (black circle) as follows: (Ci) The locations of the stimuli (red and blue circles) are chosen such that they are equidistant from the location of the mean of the system state (black circle), which implies that the stimuli have equal aesthetic values. (Cii) Similar to Ci, but with a slightly different location for blue stimuli to create an asymmetry in reward structure. (Ciii) Similar to Cii, but with a slightly different location of the red stimulus (instead of the blue one). **(Di-Diii)** Similar to Bi-Biii, but with asymmetry in reward structure induced with aesthetic reward.

We then investigate the perceptual privilege attributed to the aesthetic value of sensory input. For instance, it has been shown that aesthetic aspects of an image can bias the perceptual preference in binocular rivalry and other related paradigms (Huang et al., 2013; Mo et al., 2016; Shang et al., 2020; Tsikandilakis et al., 2019). To replicate the key findings of this category of studies (i.e., longer dominance of aesthetically preferred images), we chose the parameters of the aesthetic component of our model, such that, the aesthetic value of one image is higher than the other one. We did this by slightly changing one of the coordinates of one of the stimuli, altering the distance of stimulus from the mean of the system state distribution (turning the symmetric case ***Figure 5Ci*** into the asymmetric Cii and Ciii). Here again, we have equi-dominance for the symmetric case (***Figure 5Di***), and biased dominance durations for asymmetric reward structures (***Figure 5Dii-Diii***), similar to external reward (***Figure 5A-B***).

Through asymmetries in the reward function and the dynamics of the reward function, we can also explain all Levelt (1965)’s propositions, that state how perception during binocular rivalry depends on the attributes of the presented stimuli. Levelt’s propositions state (adapted from, Bras-camp, Klink, et al., 2015):

I. Increasing stimulus strength for one eye will increase the perceptual predominance of that eye’s stimulus.
II. Increasing the difference in stimulus strength between the two eyes will primarily act to increase the average duration of perceptual dominance of the stronger stimulus.
III. Any manipulation that increases the difference between the two eyes’ stimulus strengths will reduce the perceptual alternation rate.
IV. Increasing the stimulus strength in both eyes while keeping it equal between eyes will generally increase the perceptual alternation rate, but this effect may reverse at near-threshold stimulus strengths.

The key to compatibility with Levelt’s propositions is having the Helmholtzian notion of sensation (Dayan, 1998; Brascamp, Sterzer, et al., 2018) in our model. The Helmholtzian character of our reward function comes from two sources. One is as part of the aesthetic value (weighted like-lihood of the input, or fluency component of the aesthetic reward, see section Model of aesthetic value). The other, is as the agent’s estimation of the unlikelihood of its sensory input (for a more formal treatment, see Methods section). The former contributes as a reward for the portion of the sensory input that the agent can explain, and the latter, as the penalty, for the portion that the agent cannot explain (the mismatch between percept and actual sensory input).

In conventional, symmetric settings for binocular rivalry, the Helmholtzian contribution from unlikelihood estimation may not play a significant role. If we assume in a typical setting of binocular rivalry, sensory evidence provided by either of the presented stimuli is equally unlikely (e.g., they are equally noisy, dim, etc), then any of the possible interpretations of the sensory input leave almost half of the sensory input unexplained. Thus, in such settings, irrespective of the perceptual selection of the agent, the agent’s estimation of the unlikelihood of its sensory inputs remains almost constant over time, and thus, it may not play a significant role in shaping the fine-grained temporal dynamics of switching. This is not the case for the Helmholtzian contribution attributed to aesthetic value.

In the majority of Levelt’s propositions (except the last one), an asymmetry in the strength of the evidence for the stimuli is the central condition of the proposition (e.g., lower or higher contrast of one of the stimuli). Thus, by mapping the asymmetries in Levelt’s propositions (i.e., asymmetries in the attribute of stimuli) to the unlikelihood term of our reward function (***Equation 3***) we can readily explain these three propositions.

Corresponding settings to the first and second propositions are quite similar to the simple reward asymmetry discussed earlier (***Figure 5***), however instead of the task reward, the source of privilege is stronger evidence on one percept (the unlikelihood term) that lead to an excess of overall reward magnitude of one percept (or more precisely, more penalty for worse explanation of the weaker input). This leads to the first dominance (predominance) of the privileged stimulus (for proposition I), and overall longer dominance of the privileged stimulus (for proposition II), similar to ***Figure 5 A-B***. Our model is also compatible with the more delicate aspects of proposition II, which emphasizes the *difference* between the strength of the left and right stimuli. In our model, the difference between the expected utilities is also central (given the expected utilities of percepts are compared to each other). Thus, any changes in the strength of stimuli that lead to an asymmetry, result in a difference in the utility of left and right perception and, consequently, an asymmetric dominance duration.

The third proposition can also be explained similarly based on induced reward asymmetries that are also compatible with previous accounts (e.g., Leptourgos et al., 2020). Previous accounts of the third proposition relied on neural adaptation (Brascamp, Klink, et al., 2015). As we discussed earlier (see ***Figure 3*** and ***Figure 4***), adaptation is also present in our framework. Through this, manipulations that increase the difference between the strengths of the two stimuli, arrange for the value of one percept to remain higher than that of the other for a longer period of time (as compared to the case where both have equal strength and decay together, see ***Figure 4E***).

We can explain the last proposition (IV) based on the potential effect of the stimulus strength on aesthetic value and its temporal dynamics. We implicitly assumed that the utility stemming from basic aspects of sensory input (e.g., contrast) is part of the classical Bayesian loss function (our Helmholtzian unlikelihood components, ***Equation 3***), however, it can also be relevant in terms of aesthetic judgments (see, e. g., Tinio et al., 2011), and as a consequence, can also affect the temporal dynamics of perception. Thus, we refine our previous assumption and propose stronger stimuli (e.g., images with higher contrast), at least for a range of values have higher aesthetic values (Tinio et al., 2011).

In the model of aesthetic value that we employed, we can change the magnitude of aesthetic value by changing the geometry of the model (e.g., changing the coordinates of the stimuli in the feature space). Here, we need only change the distance between rivalling stimuli and the system state’s mean (i.e., making red and blue circles in ***Figure 4A***, closer or further from the black circle, as we also used in ***Figure 5C***). As a consequence, a change in the aesthetic value (due to distance) also leads to a change in the rate of decay of the aesthetic value, i.e., the higher the aesthetic value, the greater the decay rate. When both stimuli have the same strength (as stated in the fourth proposition), a faster decay of the aesthetic value of the perceived (dominant) stimulus also implies a faster growth of the (counterfactual) aesthetic reward of the suppressed percept (***Figure 4E***). Similarly, when the agent is exposed to weak stimuli in both eyes, the value of the dominant percept decays slowly, and the counterfactual value of the suppressed percept grows proportionally sluggishly. These together lead to faster (or slower) alternation when the strength of stimuli on both sides is higher (or weaker). The contribution of the suppressed precept (through counterfactual reward) could be even more important, given that abbreviated periods of suppression have been more emphasised in previous studies (see, e. g., Blake et al., 2002). For near-threshold stimulus strengths where a reversal has been observed (i.e., Brascamp, Klink, et al., 2015, a decrease rather increase in alternation rate was observed), other factors may contribute more (e.g., changes in the level of exploration due to low belief value in a POMDP or other aspects of inference processes) and need to be investigated further. However, it should be noted that the reversal was not observed in all cases (e.g., changing dot luminance contrast did not elicit a reversal Brascamp, Klink, et al., 2015).

### Rich temporal dynamics of binocular rivalry

As noted earlier, perceptual multistability exhibits rich temporal dynamics (see, Suzuki et al., 2007, and see ***Figure 6*** right panels) that is not completely understood. These rich dynamics in the rate of perceptual switches span a wide range of timescales: an initial increase over the course of dozens of trials (***Figure 6A***, right); a gradual decrease within a single trial (tens of seconds, ***Figure 6B***, right); a slow increases across several sessions (***Figure 6C***, right). We demonstrate, by employing a sufficiently complex reward function we are able to replicate and explain this rich temporal dynamics.

**Figure 6.**
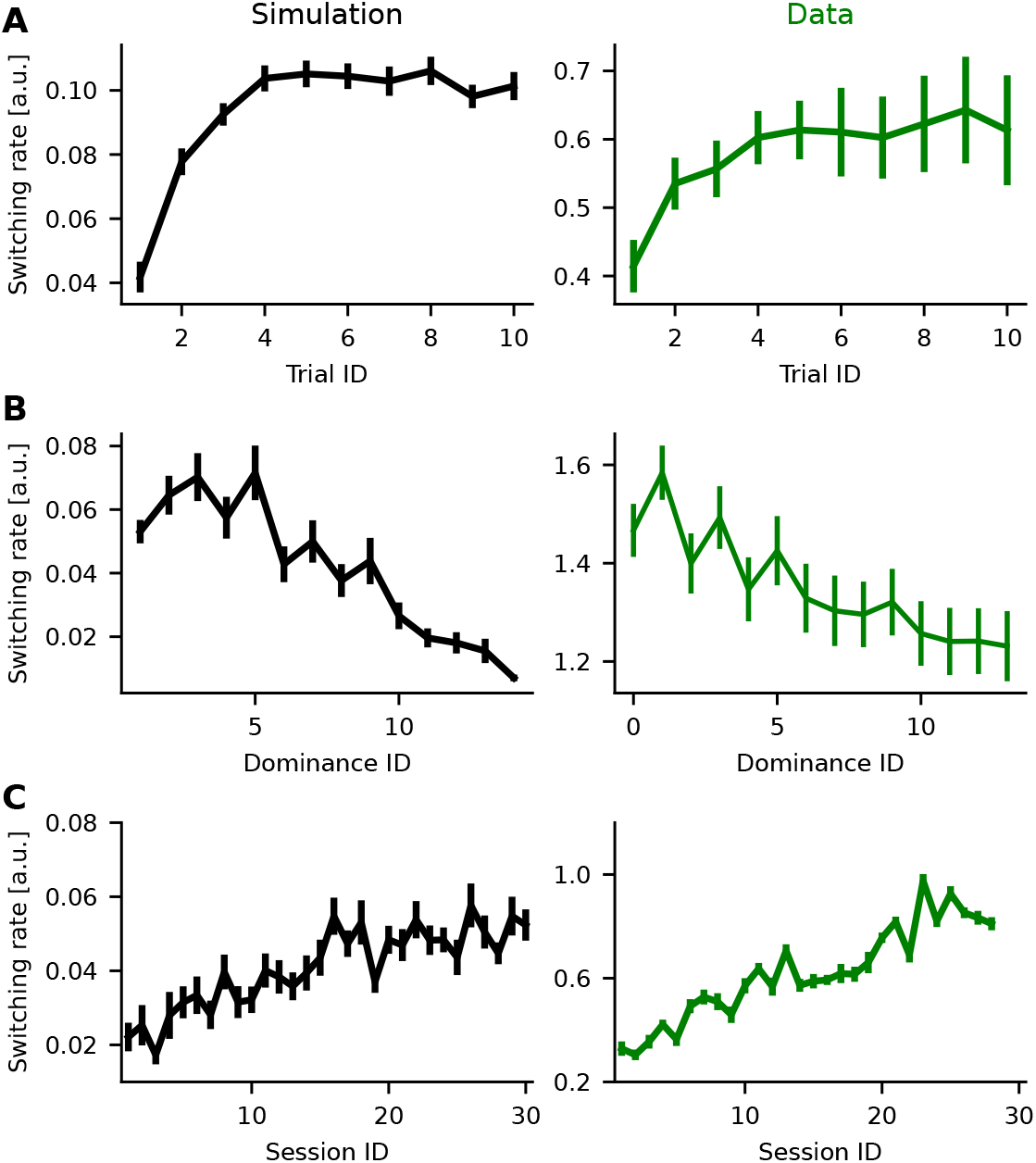
Capturing temporal dynamics of perceptual multistability at multiple timescales. **(A-C)** Modelled (left) and behavioural data of switching rates (right, from Suzuki et al., 2007) quantified as the reciprocal of the duration of perceptual dominance. **(A)** The initial increase of switching rate over 10 trials,(including all dominance periods within a trial) and averaged across multiple participants. The number of participants, varies across trials in empirical data, and it is 40 for simulation. For the data panel (right, green), switching rate were averaged across 40 participants for the first 5 trials, and 16 participants for last 5 trials (resulting in larger error bars for trials 6 through 10; see, Suzuki et al., 2007, for more details). **(B)** Decreases in switching rate across dominance instances within individual trials. **(C)** Long-term speed-up of switching rate across sessions for one participant, and a single simulated participant (individual data point for each session is average across 20 trials). The left panels are from ***Figure 1D-F***, and are presented for easier comparison with simulations (right panels). In all panels, error bars are mean ± SEM.

We suggest that the key factor shaping the initial speed-up of the switching in naive participants (***Figure 6A***) is the tendency of the agent to explore or better understand the scene (noting that Suzuki et al., 2007, only observed it in naive participants). As discussed in the Methods section, we approximate the value of information associated with a stimulus that is incompletely known (Dayan and Sejnowski, 1996) with an exponentially decreasing exploration bonus. At the beginning, stimuli enjoy a high exploration bonus; but this decays as the agent perceives them, and so gets to know them better. This implies that the rate of perceptual switches is initially low (the agent experiences longer dominance periods) and gradually increases when the exploration bonus of both stimuli is equally low. This is captured by our simulation (***Figure 6A*** left) as reported by Suzuki et al. (2007, ***Figure 6A*** right). There are also more principled approaches to potentially explain these dynamics, for instance, based on richer hierarchical inference for novel stimuli that are further discussed in section Limitations and future extensions (also see our more elaborate description of the state, section Thick state).

We suggest the gradual slow-down in the rate of perceptual switches is governed by the way that the dynamics of value evolves within the timescale of a single trial (tens of seconds). As discussed above (***Figure 4E-F***), the push-pull dynamics induced by the aesthetic value shapes the dominance periods and switching times. Along the same lines, but across several switches, aesthetic value can have a characteristic dynamic that shapes the length of dominance periods and timing of the switches in the course of a trial (***Figure 4E-F***). At the beginning of the trial, switches are fast due to the strength of the push-pull mechanism. But aesthetic value also globally decays (***Figure 4E***). This implies that the push-pull mechanism gradually loses its strength and the switches primarily are governed by noise. This is also well captured by our model (***Figure 6B*** left) as was observed empirically (***Figure 6B*** right).

For the longest timescale of perceptual dynamics, we presume that extensive perception of stimuli over several sessions (across days), increases the familiarity of the stimuli for the agent with the effect of reducing the switching cost. To be concrete, we made this be subject to exponential decay over multiple sessions. This heuristic leads to an increase in the tendency of the agent to switch and so captures the slow dynamics of the switching rate (***Figure 6B*** left). Suzuki et al. (2007) observed a large variability in the characteristics of the acceleration of the switching rate between the participants. We speculate such individual differences can be attributed to the agent’s parameter settings. For instance, a speed-up in switching rate with a larger dynamic range can be reproduced by a lower initial value of the switching cost (Supplementary Figure 2).

Overall, these results demonstrate that a sufficiently complex reward function can also capture the rich temporal dynamics of perception during binocular rivalry, and offer an explanation for the underlying computational mechanism. This implies a novel perspective on perceptual switches, where transitions from one percept to other ones are affected by a variety of cognitive factors in addition to basic bottom-up sensory processes (such as perceptual modulation due to contrast). Furthermore, these cognitive factors not only concern the attributes of the dominant percept (which has been the main focus in most previous studies), but also the suppressed percept.

## Discussion

Our account of perceptual multistability which is grounded in value-based decision-making, provides a more comprehensive treatment of the computational facets of this phenomenon and explains a diverse range of behavioural data in a single framework. Furthermore, this decision-theoretic approach addresses several shortcomings of the more traditional, purely probabilistic, Helmholtzian accounts (Brascamp, Sterzer, et al., 2018). These shortcomings encompass both behavioural and neural aspects of perceptual multistability.

Behaviorally, our computational model offers a unified explanation for a wide range of multistability data. This ranges from classical characteristics of perceptual multistability that have been known for decades, such as the distribution of the dominance periods (Blake et al., 2002), and Levelt’s propositions (Levelt, 1965; Brascamp, Klink, et al., 2015), to recent, data such as reward modulation (Balcetis et al., 2012; Wilbertz, van Slooten, et al., 2014; Marx et al., 2015; Wilbertz, van Kemenade, et al., 2017; Pothast et al., 2024; Lunghi et al., 2023; Lunghi et al., 2019; Haas, 2021; Haas, 2021) and more delicate aspects such as multi-timescale dynamics of perception (Suzuki et al., 2007; Weilnhammer, Fritsch, et al., 2021a). Thus, our model not only accounts for the key behavioural features explained by previous computational (Dayan, 1998; Schrater et al., 2007; Sundareswara et al., 2008; Hohwy et al., 2008a; Reichert et al., 2011; Moreno-Bote et al., 2011; Gershman et al., 2014; Weilnhammer et al., 2017; Leptourgos et al., 2020; Cao et al., 2021) and mathematical models (Braun et al., 2010; Deco et al., 2013; Theodoni, 2014; Whyte et al., 2023), but also extending beyond them, to elucidate cognitively richer aspects. This is particularly important in the context of recent data as a growing body of evidence suggests that perceptual dominance and perceptual dynamics can be modulated by a wide range of cognitive factors (Scocchia et al., 2014; Martin et al., 2021; Safavi and Dayan, 2022; He, 2023; Fleming et al., 2024). Neurally, several neural systems beyond sensory areas that are implicated in general aspects of decision-making (e.g., ACC, PFC) are also involved in perceptual multistability. Crucially, the computational side of our framework offers an explanation for the elusive behavioural aspects, and we suggest that its natural implementation in the brain will account for the rich collection of neural findings (see section Implication for the underlying neural substrate).

### Implication for the underlying neural substrate

Although areas of the brain are far from singular in their computational roles, it remains true that if percepts arise as choices, then regions participating in the processes of decision-making should also be engaged in multistability. This is true for the ACC (Lyu et al., 2022; Drew et al., 2021; GelbardSagiv et al., 2018), striatum (Baker et al., 2015; Sekutowicz et al., 2016), LC (at least using pupil size as proxy of its activation, Turi et al., 2018; Einhauser et al., 2008; Einhäuser et al., 2008; Naber et al., 2011; Schütz et al., 2018), PFC (see, e. g., Zaretskaya and Narinyan, 2014; Safavi, Kapoor, et al., 2014; Tsuchiya, Wilke, et al., 2015; Odegaard et al., 2017; Weilnhammer, Fritsch, et al., 2021b; Block, 2019; Michel, 2022; Kapoor et al., 2020; Dwarakanath et al., 2020) amygdala (Kreiman et al., 2002; Pasley et al., 2004), parietal cortex (Kanai et al., 2010; Zaretskaya, Thielscher, et al., 2010; Bahmani et al., 2019), pulvinar (Wilke et al., 2009; Qian et al., 2023) and various neuromodulators that were examined through pharmacology: serotonin (Carter et al., 2005), acetylcholine (Sheynin et al., 2020) catecholamine (Pfeffer et al., 2018).

We can speculate about how these various components are marshalled. For instance, the cascade leading to a perceptual switch might ultimately be triggered by the LC, noting early signs in pupil dilation (Einhauser et al., 2008) and the evidence that pharmacological manipulation of catecholamine systems increases the switching rate (Pfeffer et al., 2018). One computational correlate of this is the substantial uncertainty involved, particularly in binocular rivalry, as almost half the sensory input typically remains unexplained given either perceptual decision. Norepinephrine (NE), in concert with acetylcholine (ACh), has been suggested they reflect aspects of uncertainty, and occasioning the resetting of brain states over a large scale (Yu et al., 2005; Bouret et al., 2005). There are potential mechanistic substrates for aspects of their action (e. g., a modulation of excitation-inhibition balance by cholinergic and catecholamine systems, as suggestd by Sheynin et al., 2020; Pfeffer et al., 2018), although the exact circuits involved are as yet unclear.

Next, we suggest that the ACC (which exerts an important influence over the LC) may be involved in reevaluating perceptual actions, just as for other actions (Fouragnan et al., 2019; Brockett et al., 2020; Holroyd et al., 2021b; Akam et al., 2021). For instance, Fouragnan et al. (2019) combined fMRI, and transcranial ultrasound stimulation, in the context of a 3-arm restless bandit task to show that the values of options that are not currently available, but may potentially be chosen in the future, are represented in ACC. In their experiment, only 2 of the 3 bandits were available for choice, but the value of the third bandit still needed to be tracked. The authors showed that the ACC is critically involved in translating the value of such unavailable choices into a *counterfactual plan* that can be executed in the future. This has an obvious resonance with rivalry, with the key difference that, here, choices are a set of internal actions. Furthermore, the involvement of ACC in perceptual multistability is supported by a variety of experimental modalities (Lumer et al., 1998; Gelbard-Sagiv et al., 2018; Drew et al., 2021; Lyu et al., 2022). These data not only support the engagement of ACC in perceptual switches but also the processing of the suppressed percept (Levinson et al., 2021), which would be necessary for computing the counterfactual plan.

Equally, we suggest that sub-regions of the PFC such as ventrolateral PFC (vlPFC) execute the perceptual actions, and thus influence the timing of perceptual switches. There is growing evidence of the involvement of various parts of the PFC in perceptual multistability (Zaretskaya and Narinyan, 2014; Safavi, Kapoor, et al., 2014; Tsuchiya, Wilke, et al., 2015; Odegaard et al., 2017; Block, 2019; Michel, 2022). A particular role has been suggested for vlPFC, a region involved in cognitive control, reflexive reorienting, motor inhibition, and action updating (Levy et al., 2011). Neural responses in this region reflect the value of visual objects (Ghazizadeh et al., 2021), are selective for sensory features in conventional visual paradigms (Pigarev et al., 1979; Ó Scalaidhe et al., 1997; Hussar et al., 2009; Safavi, Dwarakanath, et al., 2018), represent perceptual content during binocular rivalry (Kapoor et al., 2020), and exhibit characteristic network dynamics in the vicinity of perceptual transitions (Safavi, Logothetis, et al., 2021; Safavi, Panagiotaropoulos, et al., 2023; Dwarakanath et al., 2020).

Although there is substantial supporting evidence, other studies suggest perceptual switches per se do not heavily involve the fronto-parietal network (see, e. g., Brascamp, Blake, et al., 2015), which is believed to be intimately involved in the sort of cognitive control that we have suggested should be engaged in multistability (e. g., Zanto et al., 2013). Most of these studies shape their questions and experiments in the context of the recent debate on the role of the frontal lobe as a neural correlate of consciousness (NCC) and visual awareness. However, our focus here is mostly orthogonal, i. e., the role of value in determining dominance and switching. Even in that context, the involvement of the frontal-parietal network has been suggested to be intimately related to the *planning* of motor actions (without execution, Brascamp, Blake, et al., 2015) which can be considered as being compatible with our *goal-centered* approach to perceptual multistability (also inline with other proposals, see, e.g., Leopold et al., 1999; Jensen et al., 2023).

Finally, neuromodulators other than norepinephrine and acetylcholine, which play a critical role in shaping distributed information processing (Dayan, 2012), and are deeply involved in many psychiatric disorders (Iglesias et al., 2017), are also involved in perceptual multistablity (Carter et al., 2005; Sheynin et al., 2020; Pfeffer et al., 2018; Novicky et al., 2022), albeit in ways that are not completely clear. It is tempting to speculate that dopamine might be involved in at least some aspects of the complex temporal dynamics of the rate of switching (van Ee, 2005; Mamassian et al., 2005; Suzuki et al., 2007; Brascamp, Knapen, et al., 2008; Klink, Brascamp, et al., 2010). The basis for this is the suggestion that tonic levels of dopamine, by representing the local average reward rate, act as an opportunity cost for the passage of time, and thereby influence the vigour or alacrity of action (Niv et al., 2007; Mazzoni et al., 2007; Shadmehr et al., 2019). Intuitively, if the average rate of reward is high, every unit of time in which a reward is not delivered is costly for the agent, thus, it is worth the agent making decisions faster even if the costs of doing so (e. g., perceptual switches) are greater. The converse is true if the average rate of reward is low (Niv et al., 2007).

### Limitations and future extensions

This is only the first formal realization of our theory, and so suffers from various limitations. First, we assume that there are only two plausible candidates for the state of the world. The space of candidates should, in principle, be considerably larger (in keeping with the potential richness of our perceptual experiences). Furthermore, in binocular rivalry, in addition to exclusive [dominant] percepts, humans (and perhaps also non-human primates) also experience, piecemeal, superposition (see, e. g., Klink, Brascamp, et al., 2010; Skerswetat et al., 2023) and fused percepts (for special stimuli like rivalling faces, Klink, Boucherie, et al., 2017, participants perceive morphed faces that slowly switch). In fact, the factors affecting piecemeal perception may shed light on the cognitive aspects of multistability. For instance, participants who are more open to exploration (based on measures of personality, Antinori et al., 2017), exhibit longer durations of piecemeal percepts. Thus, a pressing direction is to allow and investigate a richer state space.

Second, we assumed that the agent’s observation function (***Equation 1***) has a known mean, however, the mean of this distribution should also be estimated by the agent (for instance, using a similar procedure as Khalvati et al., 2021). The resulting decreasing exploration, which we characterized using a simplistic heuristic, might then contribute to the temporal dynamics of perception in early phases, and would interact with expectations about subsequent change.

Third, in our formulation, we consider the consequences of the agent’s action to be both making a decision that the state is as implied by the associated percept, and also lowering observation noise about it (compared to the suppressed percept) which can be considered as a form of attention (Dayan and Solomon, 2010; Chebolu et al., 2022). The latter operation is computationally richer in hierarchical accounts of perception (Dayan, Hinton, et al., 1995; Dayan, 1998; Lee et al., 2003; Hohwy et al., 2008b). In such accounts, uncertainty about the unperceived image will be progressively greater at higher levels in the hierarchy, in a way that might influence the temporal dynamics. Given the explicit computational account of attention in our model, this could also provide an insight, if and how the underlying computational operation is related to the still controversial (Maier et al., 2021) question in multistability about the potential role of attention, a construct that itself is enduringly complex (Wu, 2023; Heeger, 2017; Li et al., 2017).

Fourth, our approximate POMDP solution only takes into account the immediate horizon; however, each action should really be taken based not only on its immediate return, but also on its long-term ramifications. The effects of this on the aesthetic value component of reward would be particularly complicated since this component is dynamic itself. However, some facets of perceptual dynamics that we explained based on heuristics (e.g., exploration bonus) might derive naturally through a more presbyopic solution of the POMDP.

Fifth, there are delicate aspects of the temporal dynamics of perception (such as the unit of time and variability of dominance periods across participants) that merit further investigation. For instance, the unit of time in the current setting of the model is a free parameter, so the model’s temporal dynamics follow the experimental data (***Figure 6***), but only up to an arbitrary scaling factor. It would be interesting to relate the actual timescales in the model, such as the exponential decay and the learning rate of the system state, to empirical observations of switching behaviour. Furthermore, it is also interesting to investigate whether and how this timescale is related to the characteristic timescale of other cognitive functions, and, indeed, whether it is influenced by dopaminergic neuromodulation, given the latter’s effect on other aspects of timing behaviour (Meck, 1983).

Finally, although the incorporation of aesthetic value (Brielmann and Dayan, 2022) was a key step towards a novel approach to sensory processing in multistability, the full potential of the aesthetic model was not explored in this study. Upon exposure of the agent to a new sensory object, the construction of a percept likely require some initial computation to stabilize the corresponding percept (an initial inference process). As a consequence, the value of the sensory object is not known to the agent. Given in our model, the entire inference process is boiled down to a simple Bayesian belief update (see Partially observable Markov decision process), the aesthetic model can be based on a *hierarchical* percept formation that requires some time to be realized. In the current formulation, for simplicity, we use a heuristic (***Equation 5***) to incorporate this initial computation. However, this can be incorporated more naturally into the model, for instance, as a richer hierarchical inference process in the POMDP (also related to the third point discussed above) or in the aesthetic model through the initial dynamics of the system state as a consequence of an initial inference process (also see our more elaborate description of the state, section Thick state). Moreover, the choice of parameters for the aesthetic component of the model was limited to a form of decay. This was motivated by the simple choices of visual images (e.g., gratings) that were used in a large body of previous experiments. However, the few studies that also looked at the temporal dynamics induced by more complex stimuli (like faces) reported non-monotonic temporal dynamics (see, e. g., Klink, Brascamp, et al., 2010; Klink, Boucherie, et al., 2017). The model of aesthetic reward can also have a nonmonotonic temporal structure, under more complex parameter settings (see, e. g., Brielmann and Dayan, 2022; Brielmann, Berentelg, et al., 2023). There is also room for further expanding the aesthetic model to better explain the temporal dynamic of multistablity. For instance, to explain the long-term speed-up of switching (***Figure 6***), we introduced a heuristic (gradual change of switching cost across sessions). Such slow dynamics could be potentially explained through incorporating a slow learning process that affects the coordinate of the expected true distribution in the feature space of the aesthetic model (the expected true distribution was assumed to be constant in the original model, Brielmann and Dayan, 2022). Presumably, such slow learning should happen through offline processes, which also matches the slow dynamic in sessions distributed across days. Thus, it is pressing to investigate if and how more complex temporal dynamics of perception in binocular rivalry are shaped by the temporal dynamics of the aesthetic value.

### Prospects

Perceptual multistability, which has been an object of study for many decades, arises in a wide range of species, across almost all sensory modalities, and is also relevant for a wide range of neural computations (Safavi and Dayan, 2022). It therefore has the potential to provide a unique window for understanding a wide range of cognitive functions and dysfunctions. In the following, we discuss some of the opportunities our novel perspective opens up.

First, having a handle on the temporal dynamics of multistability, may provide new understanding of distinctions between healthy and psychiatric populations (see, e. g., Ye et al., 2019; Jia et al., 2020), as most reported differences concern these facets. For instance, we speculate that, in particular forms of depression, value estimates will be biased lower, explaining the slower switching rate; and in anxiety, the informational value may have a higher weight (particularly for individually-scary stimuli), leading to elevation in the rate of perceptual switches (Jia et al., 2020). Equally, conditions in which dopamine release or reception differ (including forms of schizophrenia and depression; Maia et al., 2017; Pizzagalli et al., 2019) may exhibit different switching rates, given the putative tie to vigour and timing mentioned above (Niv et al., 2007; Meck, 1983). Altered rates of perceptual switches have also been reported with animal models of psychiatric conditions, including mouse models of autism (Palagina et al., 2017; Bogatova et al., 2023) (also in keeping with human studies; Robertson et al., 2013; Spiegel et al., 2019). The recent development of no-report paradigms (Tsuchiya, Wilke, et al., 2015), makes multistability an appealing paradigm for studies with animals and even children (Hudak et al., 2011; Beers et al., 2014; Karaminis et al., 2017).

Second, our formulation of perceptual multistability offers a functional and computational interpretation for some traditional biophysical characteristics such as adaptation. Besides the Helmholtizan approaches (Brascamp, Sterzer, et al., 2018), a large body of previous mathematical approaches (e.g., work on circuit and synaptic principles such as attractor dynamics, adaptation, and mutual inhibition; Braun et al., 2010; Deco et al., 2013; Theodoni, 2014; Whyte et al., 2023) could account for many characteristics of perceptual multistability (e. g., distribution of dominance periods). However, they were largely agnostic to the sort of computational purposes on which we have focused. For instance, as we discussed earlier (see ***Figure 3*** and ***Figure 4***), we introduce adaptation in our framework in the cognitive context of boredom. Such formulations tie together biophysical and cognitive constructs (Scocchia et al., 2014; Safavi and Dayan, 2022). This is in keeping with other recent data – for instance, traditional visual adaptation (Keller et al., 2017) and pupil modulation by brightness (Turi et al., 2018) has been shown to be controlled by other cognitive processes. Future studies should address how such biophysical principles can be fully integrated with our normative computational framework.

Third, our approach suggests a link between modern conceptions of internal reinforcement learning (Hazy et al., 2007; Dayan, 2012; Lieder et al., 2018), sensory processing, and even neurofeedback (Lubianiker et al., 2022). Thus, it would be particularly important to examine cognitive control in this context (also given other suggestions, see, e. g., Botvinick et al., 2001; Drew et al., 2021). Indeed, the three components of cognitive control (monitoring, specification, and regulation) proposed by Shenhav et al. (2013), are also applicable to perceptual multistability. Monitoring concerns the assessment of the current circumstances and the evaluation of the performance of the ongoing strategy. For perceptual multistability, this means (self-)assessing the quality of inference about the state of the world (which is the primary role of perception). Specification deals with the choice of actions or sub-tasks, that in the case of multistability are, for instance, the exclusive perception of one of the stimuli, or a perceptual switch. Finally, regulation concerns the adjustment of lower-level information processing machinery. In terms of perceptual multistability, this would be critical when elements of the task, or the statistics of the world, change. At a mechanistic level, adjustments could involve several processes, for instance, adaptation and tuning of the excitation-inhibition balance, possibly with computational implications (e.g., see the earlier discussion on adaptation).

Finally, since our proposed computational framework (and indeed the aesthetics model that we suggest endows percepts with value; Brielmann and Dayan, 2022) exploits Helmholtzian notions of sensation as inverse generation, our results are consistent with many others in that tradition (Brascamp, Sterzer, et al., 2018; Safavi and Dayan, 2022), and can readily be extended to incor-porate the developments in these accounts. This would include, for instance, a more thoroughly hierarchical treatment (Dayan, Hinton, et al., 1995; Dayan, 1998; Lee et al., 2003).

In sum, in this study, we have introduced a decision-theoretic perspective on perceptual multistability, using binocular rivalry as an illustration of our computational framework. This framework not only explains several elusive aspects of the phenomenon (such as reward modulation), but also opens up modern conceptions of internal reinforcement learning in the service of understanding perceptual phenomena and sensory processing in health and disease.

## Methods

### Perceptual multistability as a POMDP

#### Partially observable Markov decision process

Humans and animals (generically, agents) frequently encounter situations in which they repeatedly need to choose one action among multiple alternatives. Each action in a sequence must be made not only based on its immediate effect in terms of gaining reward, but also based on any long-term ramifications from changing the circumstances or state of the agent. A Markov decision process (MDP, Sutton et al., 2018) is a general framework for modelling a wide range of such decision processes. The four components of an MDP are: states (specifying the circumstances of the human or animal in the external world); actions (the choice between available alternatives); transition probabilities (characterising how the state of the world might change given each possible action); and rewards, which specify the loss or gain associated with the state, action, and transition. Long-run values, a measure of the expected summed future reward, are a consequence of the agent’s policy (i.e., the way it chooses to act at each state); and this policy can be optimised to maximise those long-term values.

Although MDPs have proved very useful for characterising many environments, one restrictive assumption is that the actor must know the state completely at all times (for instance, based on sensory input). This is not valid for many real-world problems, including our characterisation of perceptual multistability, where we assume that agents also make decisions about what to perceive and for how long (Martin et al., 2021; Safavi and Dayan, 2022). We therefore use a more complicated but more realistic class of models called *partially observable* MDPs or POMDPs (Littman, 2009). POMDPs are MDPs in which the agent may be uncertain about the state of the world, for instance, due to noisy sensory information. This noise is the product of an observation function that assigns probabilities to possible observations depending on the agent’s states and actions. The agent then needs to form and update a belief about the state of the world in light of observations. It can also choose actions based on the information they are expected to provide about the state, i.e., on the value of the reduction in uncertainty about the state to which they are expected to lead. The agent’s task remains to determine a policy that maximises long-term values. In the following, we describe the key components of the POMDP in relation to perceptual multistability, and leave more generic aspects and procedures for the Appendix (see section Partially observable Markov decision process).

#### Sensory, perceptual and decision processes

As in conventional Bayesian accounts of perception, we assume that our decision-making agent has a generative model of the world (Hinton et al., 1997; Dayan, 1997; Rao et al., 1999; Doya et al., 2007; Yuille et al., 2006). which specifies what input activity would be generated given a particular state of the world. Perception is then the recognition task that inverts the generative model, i.e., mapping the activities in the sensory system to the percept (or strictly, a distribution over possible percepts). We consider there to be a single actual state *s*_*t*_ ∈ 𝒮 of the world at any one time *t*, that must be inferred from sensory observations. However, rather than sticking to the regular probabilistic recognition model (inverse of the generative model, Dayan, 1997), we consider the fuller body of Bayesian decision theory for recognition. This is a key difference between this model, and previous models of perceptual multistability with the same spirit (Dayan, 1997; Gershman et al., 2014).

We build our account in two stages: first, assuming a very simple generative model containing only two possible states (see Thin state); and second, considering that the states have extra dimensions of internal structure (see Thick state) that need to be inferred over time from sensory input in a way that can benefit from the allocation of attention (which we treat itself as another component of the action space for the POMDP).

#### Thin state

For the simplest formulation of binocular rivalry, we assume there are only two plausible, but incompatible candidates for this state, 𝒮 = {+1, −1} (Dayan, 1998). Thus, the agent should make an inference about the actual state at time *t* – this is the perceptual decision 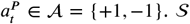. 𝒮 and 𝒜 are the state and perceptual action spaces of our perceptual POMDP; what remains is to specify the observations, transitions, and rewards.

In general, the observation function of a POMDP,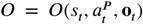, expresses the probability distribution over possible observations **o**_*t*_ made at time *t*, given that the agent took action 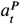 and ended up in state *s*_*t*_. In our case, the observations only depend on the current state *s*_*t*_ of the world, and not the perceptual action. We make the simplifying assumption that there is one scalar input for each eye, with Gaussian noise, so

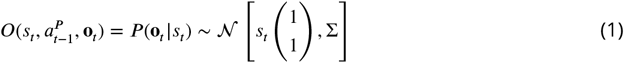

The mean of this distribution, expresses the expectation that the eyes should see the same stimulus. The covariance matrix Σ = *σ*^2^ℐ expresses the noise surrounding this stimulus; for the moment, we consider this to be proportional to the identity matrix (ℐ). Note that, in the case of binocular rivalry, a characteristic observation might express the conflict between the content of the two eyes (and the noise).

The transition function 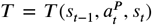 of the POMDP expresses the probability distribution over the ending state, *s*_*t*_, given that the agent started from state *s*_*t*−1_ and took action 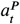. This accommodates the possibility of change in the environment, and so will be intimately tied to the dynamic nature of our perceptual experience (Pastukhov et al., 2022). Again, for our perceptual POMDP, the transition is independent of the perceptual action, leaving

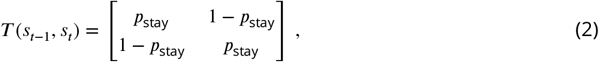

where *p*_stay_ denotes the probability of staying in the original state of the world, and we assume is known to the agent.

The reward function has a number of components: a loss function 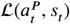 associated with a mismatch between the percept 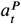 and the state *s*_*t*_ as informed by the history of observations up to time *t*, 𝒪*t* = {**o**_1_, …, **o**_*t*_}, a cost 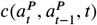 of switching from percept 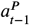 to percept 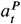 which can vary over a long time-scale, an aesthetic reward 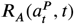, an (heuristic) exploration bonus 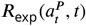, and an external reward 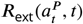 associated with what the agent can achieve the world based on the percept (as in various of the experiments described above). We cover these in turn.

The standard Bayesian decision theory articulates the loss associated with choosing action 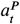 when the true state of the world is *s*_*t*_. In a filtering context, one Helmholtzian possibility for this (slightly abusing the notation) is

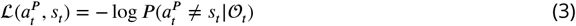

effectively counting the unlikelihood of the observations 𝒪*t* given the current assignment of state 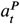. A prominent component of this is the negative log probability of the current stimulus **o**_*t*_ under 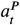

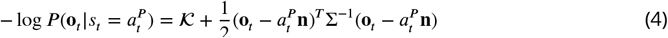

where **n** = (1, 1)^*T*^ and *𝒦* is a constant. Variants of this unlikelihood can explain perceptual preferences for higher (over lower) contrast inputs in rivalry, when the penalty for not explaining the high-contrast input is higher.

Switching between percepts (i.e., if 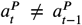) is a form of costly *internal action* (Dayan, 2012) – the equivalent of the external action of moving the eye gaze from one object to another (e.g., Callaway et al., 2021). We allow the switching cost,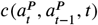, to change over a very long timescale (i.e., experimental sessions conducted across days). That is, we assume that, within each experimental session (as structured in Suzuki et al., 2007) with multiple trials (each with length *τ*_tr_, with an inter-trial interval of a few seconds), the switching cost remains constant. However, this constant switching cost decreases exponentially over multiple sessions (at least spread over days, as in Suzuki et al., 2007), for instance from an increase in the familiarity of the stimuli.

Balancing out these costs are three rewards. The aesthetic reward, 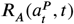, follows the model of Brielmann and Dayan (2022, as briefly described in section Model of aesthetic value in the Appendix). This theory suggests that aesthetic value is derived from optimising a sensory system to process present and future stimuli well, assuming, consistent with our Helmholtzian picture, that the agent has a *momentary or contextual* probabilistic model over the whole collection of stimuli it thinks it might sense (denoted as the system state) and a permanent probabilistic model for the sensory input it expects in the future (called the expected true distribution). Thus, the aesthetic reward is computed on the basis of current and past percepts and agent’s generative models. At every moment, the agent computes and accrues the aesthetic reward from the perceived (dominant) stimulus, but also computes a counterfactual aesthetic reward for the suppressed percept. Both consumed and counterfactual aesthetic rewards are computed as described in section Model of aesthetic value in Appendix, with the key difference that the consumed one is computed based on the location of dominant stimulus in the feature space and the counterfactual one, based on the location of the suppressed stimulus.

We also consider a heuristic exploration bonus, 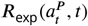. This is an approximation to the value of information associated with a stimulus that is incompletely known (Dayan and Sejnowski, 1996). This utility is attributed to all possible states, and decreases during the time that state is being perceived. With the current, binary, state space, this term is notional, and we include it for completeness; when there is information content to a state, this stands in for the benefit of learning about this information. For the moment, we consider that, as the dominance time of a given state increases, the stimulus becomes better known, at least up to forgetting, and thus less novel. We define it as:

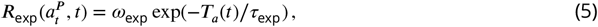

where *ω*_exp_ determines the gross contribution of this component, *T*_*a*_(*t*) is the integrated total dominance time of percept *a*, and *τ*_exp_ is the time constant according to which the bonus decreases. We assume the agent computes the integrated dominance *T*_*a*_(*t*) with a forgetting rate of *η*, given the limitation on the memory on past perceptual experience within a session and the possibility of change. Let 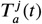 be the total dominance period of percept *a* in trial *j* up to time *t*; then we assume the cumulative dominance *T*_*a*_(*t*) (within the session) is the sum of the dominance period of the current trial and the discounted value from the previous trial (due to forgetting),

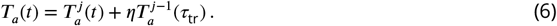

where *τ*_tr_ is the length of a trial.

Finally, the external reward 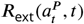 is, for our current purposes, a constant term (to be assigned to one particular percept) to account for rewards induced by the experimentalist such as associating reward to a stimulus or a percept (similar to the experimental design of previous studies, Balcetis et al., 2012; Wilbertz, van Slooten, et al., 2014; Marx et al., 2015; Wilbertz, van Kemenade, et al., 2017; Wilbertz and Sterzer, 2018) and/or rewards associated with an external, visually-guided task (as introduced in previous studies Paffen et al., 2008; Dieter et al., 2016; Moreno-Sánchez et al., 2019; Wang et al., 2021). Elements of this external reward (for instance, other remunerated decisions that can be made on the basis of the nature of the state) are rather nominal in the simplistic binary formulation. The full reward at time *t* is then

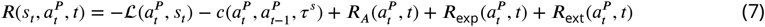

The solution of the POMDP is the optimal perceptual policy that maximizes the long-run summed expected reward (see Partially observable Markov decision process in the Appendix for more details). In the general formulation of a POMDP, both the immediate and the expected future reward (which here is dynamic, as not all the terms in the reward function are stationary) should be considered, but, to keep the model as simple as possible, we only consider the immediate reward, i.e., solving it myopically.

#### Thick state

Although the version of the model with a thin, binary, state is convenient to illustrate the workings of the model, it lacks the depth to encompass some key effects associated with the three forms of reward. In a more complete setting, we would have richer states **s**, which have internal structure that can be inferred from the observations. Here, given the state **s**_*t*_, the mean of the observations would be **s**_*t*_ ⊗ **n**. However, in rivalry, the actual input would be a noisy version of **s**^*l*^ to the left eye and **s**^*r*^ to the right eye (just as it was +1 to the left eye and −1 to the right eye in the thin state model). The agent has to make inferences about **s**_*t*_ given this conflicting input over time, and its expectations about change.

Here we ignore the possibility of mixed or averaged percepts (but see, Dayan, 1998), and only consider ocular rather than stimulus rivalry (Logothetis et al., 1996). Thus, we consider that the agent makes a composite action: one part is choosing from which of the eyes the percept will come, and, since it will then take some time to assimilate and process the information from that eye, the other part is a probabilistic description or belief about what the stimulus is, given the observations and the ocularity decisions. We do not introduce this probabilistic description formally (as it is not needed for the purposes of this simple model). However, it conceptually underpins certain aspects of the aesthetic reward. For instance, this probabilistic description of the content of the stimulus enriches and justifies the aesthetic reward, engenders an exploration bonus (since it may take some time to accumulate the relevant information), and also may provide the substrate for a suitably remunerated external decision.

One important facet of this perceptual decision-making process is that choosing that the percept is associated with the observation in one eye is to treat the components of the input from the other eye that do not match as noise. One formal possibility for this is to enrich the observation function of the generative model to allow for the possibility that the input of one eye can sometimes have excessive noise, such that

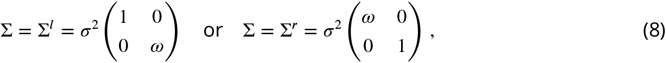

where *ω* ≫ 1 represents excess noise. Then, we can treat the ocular component of the perceptual action as also determining the version of the covariance matrix used to determine the Bayesian loss function, in the equivalent of ***Equation 4***. This would have a rather similar effect to the way that Dayan and Solomon (2010) treated attention as suppressing noise, albeit in their case from confounded receptive fields, and with a different generative model.

## Acknowledgments

This work was supported by the Max Planck Society and the Alexander von Humboldt Foundation. SS acknowledges the add-on fellowship from the Joachim Herz Foundation. We would like to thank Satoru Suzuki for generously providing the data used presented in ***Figure 1D-E*** (as well as, ***Figure 6***); Jan Brascamp for their valuable comments on an earlier version of this manuscript; scidraw.io for providing a free repository of high-quality scientific drawings (in particular Akshay Markanday for providing his “Monkey face cartoon” in this repository); and “macrovector_official” / Freepik for providing free vector graphics (icons and avatars) that were used in ***Figure 2***.

## Appendix

### Method details

#### Model of aesthetic value

We used a modern theory of aesthetic value (Brielmann and Dayan, 2022), which suggests that the utility of a sensory object derives from its influence in adapting the brain’s sensory processing system to the demands of the stimuli that it expects to encounter in the present and over the long run. As this model was introduced and described extensively in the study of Brielmann and Dayan (2022), we restrict ourselves to a brief exposition of its key aspects (and our own notation).

Here we represent stimuli, or in the context of this study, ŝ_*t*_, which is the estimate of the agent for the richer state **s**_*t*_ (introduced earlier in section Thick state). In principle, this formulation allows for different qualities of the perceptual experience through estimated stimuli. However, for simplicity, here we assumed that this estimate is limited to exact features of either left or right stimulus, i.e., we assume, for instance, there is no morph perception (e.g., as reported by Klink, Boucherie, et al., 2017) or piecemeal perception (Carmel et al., 2010).

Based on the theory of aesthetic value (Brielmann and Dayan, 2022), we assume that the agent has a *momentary or contextual* probabilistic model of what it thinks it might sense (called the system state, denoted by ***X***_*t*_), and a second model of the sensory input for the sensory input that the agent expects over the long run (called the expected true distribution, denoted by *p*^T^). Following Brielmann and Dayan (2022), we assume that both momentary and long-term distributions are normal

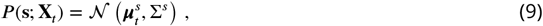

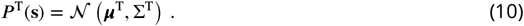

The system state is dynamic, i.e., with continuous exposure to a stimulus (perception of a stimulus), the system state gradually evolves toward the presented stimulus, thus 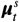 continuously adapts with a constant learning rate of *α*,

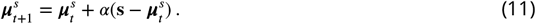

This change, relative to stimulus and the expected true distribution, determines the magnitude of aesthetic value.

The immediate perceptual reward, *r*(*t*), is defined as the log-likelihood of a stimulus given the system state at time *t*,

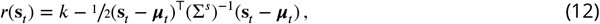

where *k* is a constant. Notably, this has a similar formulation to the one we introduced earlier in ***Equation 4***, in which we treated the the *negative* log-likelihood of the observed ocular input as a component of the loss function – and, indeed, in an hierarchical formulation of a generative model of sensory input (Dayan, Hinton, et al., 1995; Brascamp, Sterzer, et al., 2018), these terms would also be directly combined in the loss function. In the simple characterisation here in which we do not explicitly model a belief state over the perceptual details of the input, we imagine that the internal percept is exactly the input to one of the two eyes, and is directly aesthetically valued.

Stimuli that generate substantial learning or change also have a high long-term utility, thus, Brielmann and Dayan (2022) defined the second component of the aesthetic reward as follows. The system state has an inherent value that is the average expected future sensory reward, which we write here as:

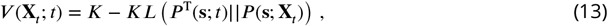

where *K* is a constant, and *KL* is the Kullback-Leibler divergence between the distribution over those stimuli implied by the momentary system state, *P* (**s**; **X**_*t*_) at time *t* and the expected true distribution *P* ^T^(**s**; *t*), that the agent believes at time *t* will arise in the long-term future (inspired by temporal difference error from reinforcement learning, Sutton et al., 2018). Thus, the utility stemming from the change in system state, Δ***V***, follows as,

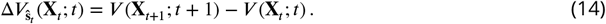

acknowledging explicitly that the change to **X**_*t*_ is occasioned by ŝ_*t*_. Ultimately, the aesthetic value, ***R***_***A***_(**s**, *t*), is defined as the weighted sum of these two components (and potentially with the constant weight, *w*_0_),

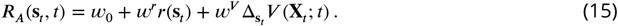

In the description of our decision-making model (section Perceptual multistability as a POMDP) we used *a*^*P*^ (the perceptual action) instead of characteristics of the state (ŝ) as a shorthand notation, thus we referred to aesthetic value by 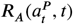.

#### Partially observable Markov decision process

##### Belief formation and maintenance

The agent forms and maintains a belief about the state of the world (as described earlier, in section Thin state), i.e., the agent’s posterior distribution over states, given its observations. Let *b*_*t*−1_(*s*) denote the probability assigned to state *s* by belief state *b*_*t*−1_ and let *b*_*t*_ be the updated belief state given the old belief state *b*_*t*−1_, action 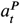, and observation *o*_*t*_. In a general form, the agent’s belief can be updated as follows,

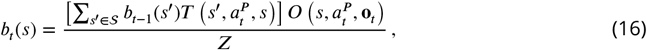

where ***Z*** is a normalizing factor (ensuring the sum of beliefs remains 1). Given that the perceptual actions of the agent do not change the state of the world, the state transition function does not depend on the perceptual actions of the agent, therefore, belief update can be slightly simplified,

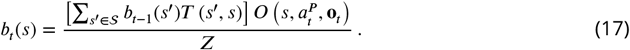

##### Solving the POMDP

The optimal decision/policy allows us to create a mapping from the belief states to the optimal actions, i.e., selecting actions that lead to the maximum utility for the agent. Based on Kaelbling et al. (1998)’s formulation, we use the notion of state estimator function, 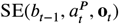, which provides an updated belief state, based on the previous belief state at time *t* − 1, the selected perceptual action and observation at time *t*:

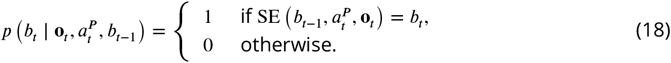

The reward function in the belief MDP is derived from the original reward function via

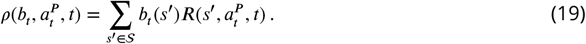

The objective is finding the optimal policy, *π*^*\**^, which in this case is only the immediate action (as we solve the horizon 1 case). Thus we need to have an estimate for the expected return for performing some action in a given belief state *b*. These are called state-action or Q-values:

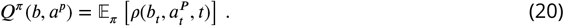

As the goal is to execute an optimal policy, *π*^*\**^, the action leading to maximum return should be selected self-consistently:

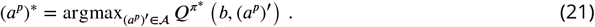

This provides a mapping from belief at time *t* to a perceptual action 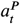.

---

**Supplementary Figure 1.**
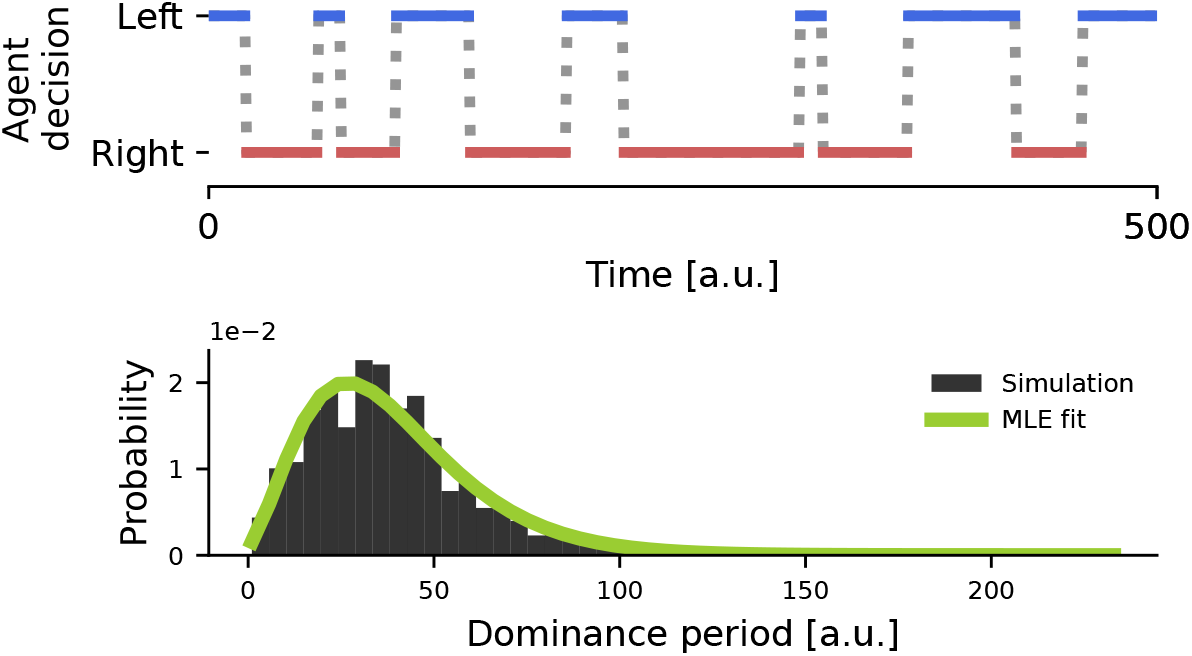
A simple POMDP with reward asymmetry. **(Top)** An example trajectory of the agent’s decision sequence between left and right images in a binocular rivalry task when one percept is rewarded (right percept is rewarded). **(Bottom)** Distribution of dominance periods for an extended length of simulation. Gamma distribution fitted with maximum likelihood estimation (MLE) and indicated with a green line.

**Supplementary Figure 2.**
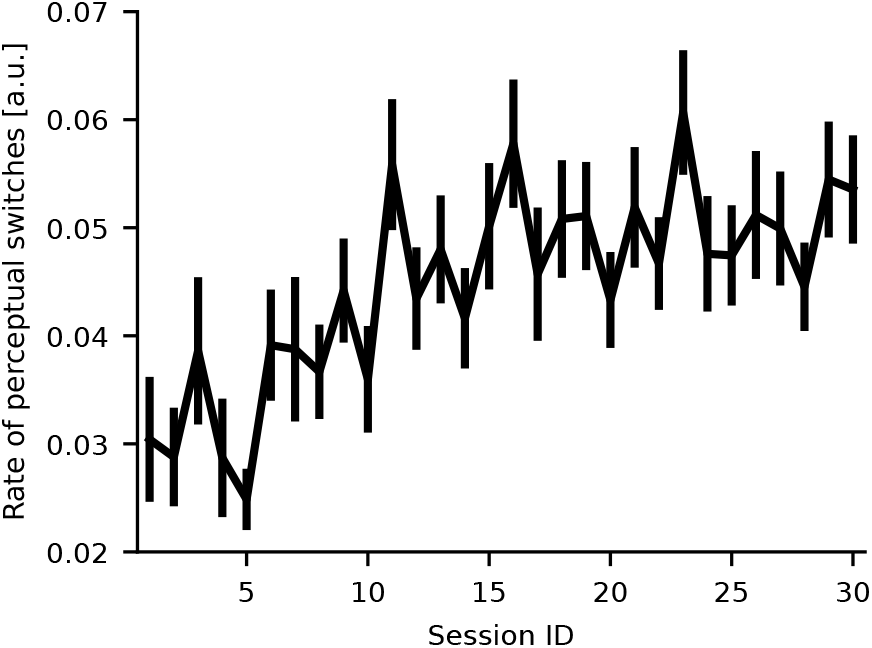
Temporal dynamics of perceptual multistability across sessions – an example. Similar to ***Figure 6***, long-term speed-up of switching rate across sessions for one participant, and a single simulated participant. Switching rates are quantified as the reciprocal of perceptual dominance durations. Individual data points for each session are averages across 5 trials.

